## Supplementary material for "A Modified BPaL Regimen for Tuberculosis Treatment replaces Linezolid with Inhaled Spectinamides": pdf

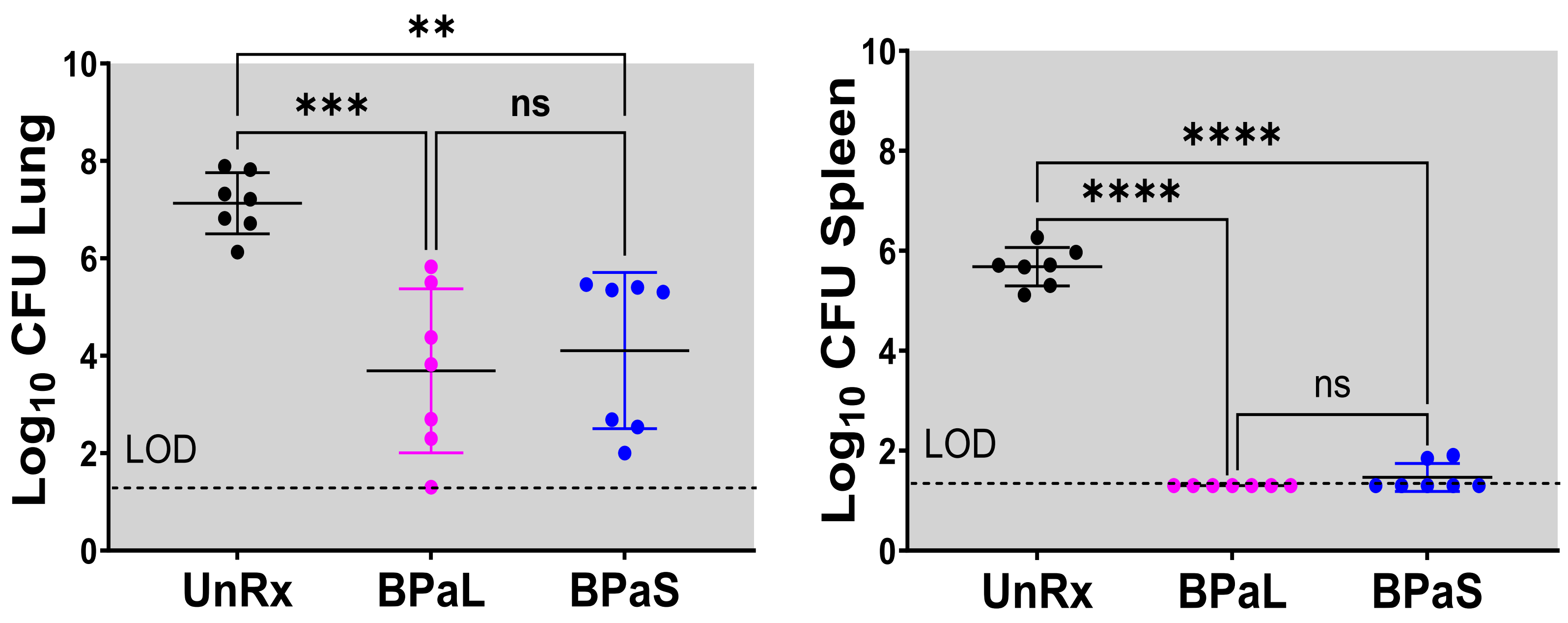


**Figure S1 (C3HeB/FeJ Study 1).** Bacterial burden (CFU) in Mtb infected C3HeB/FeJ mice treated with BPaL and BPaS regimen for 4 weeks. n = 7, LOD: limit of detection, $p < 0.05$

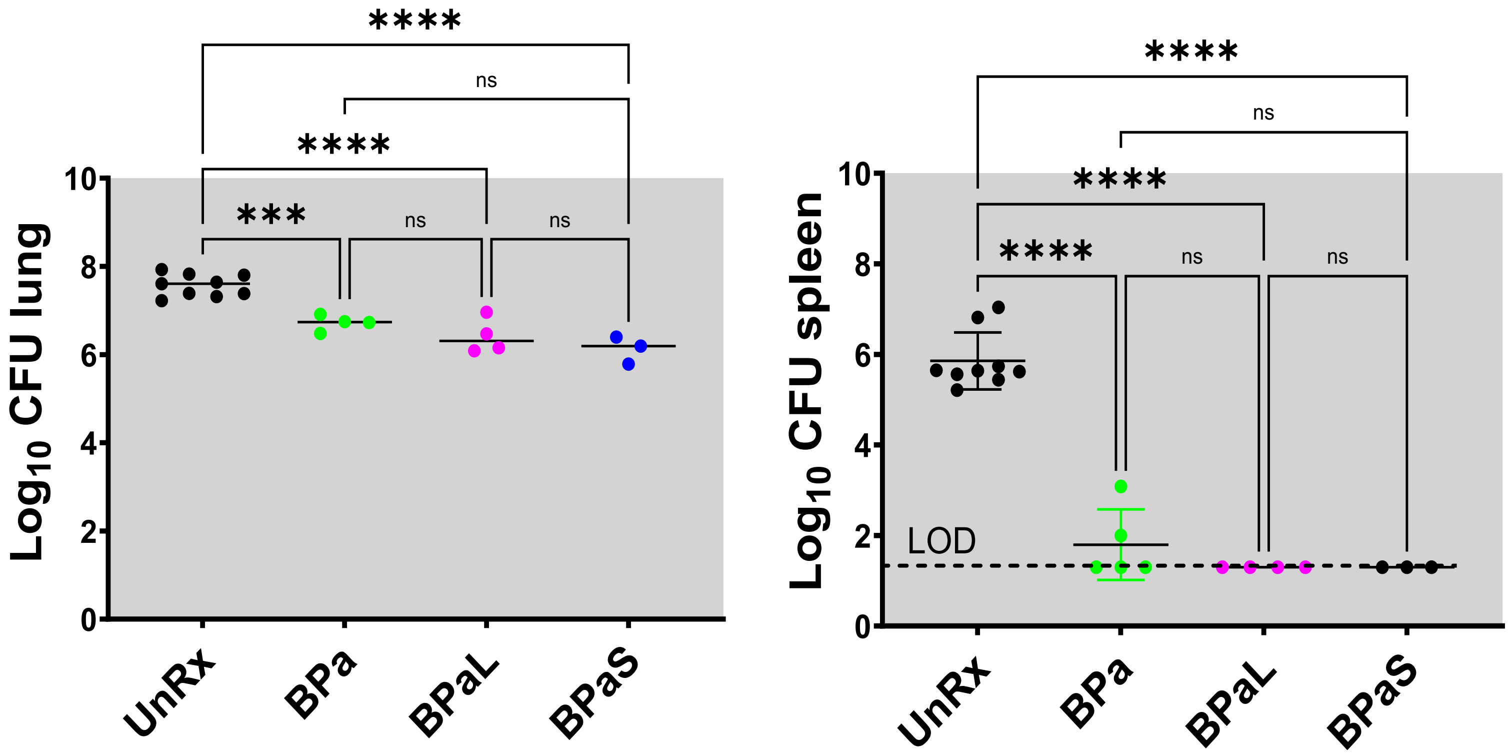


**Figure S2 (C3HeB/FeJ Study 2).** Bacterial burden (CFU) in Mtb infected C3HeB/FeJ mice treated with BPa, BPaL and BPaS regimen for 4 weeks. n = 3-9, LOD: limit of detection, $p < 0.05$

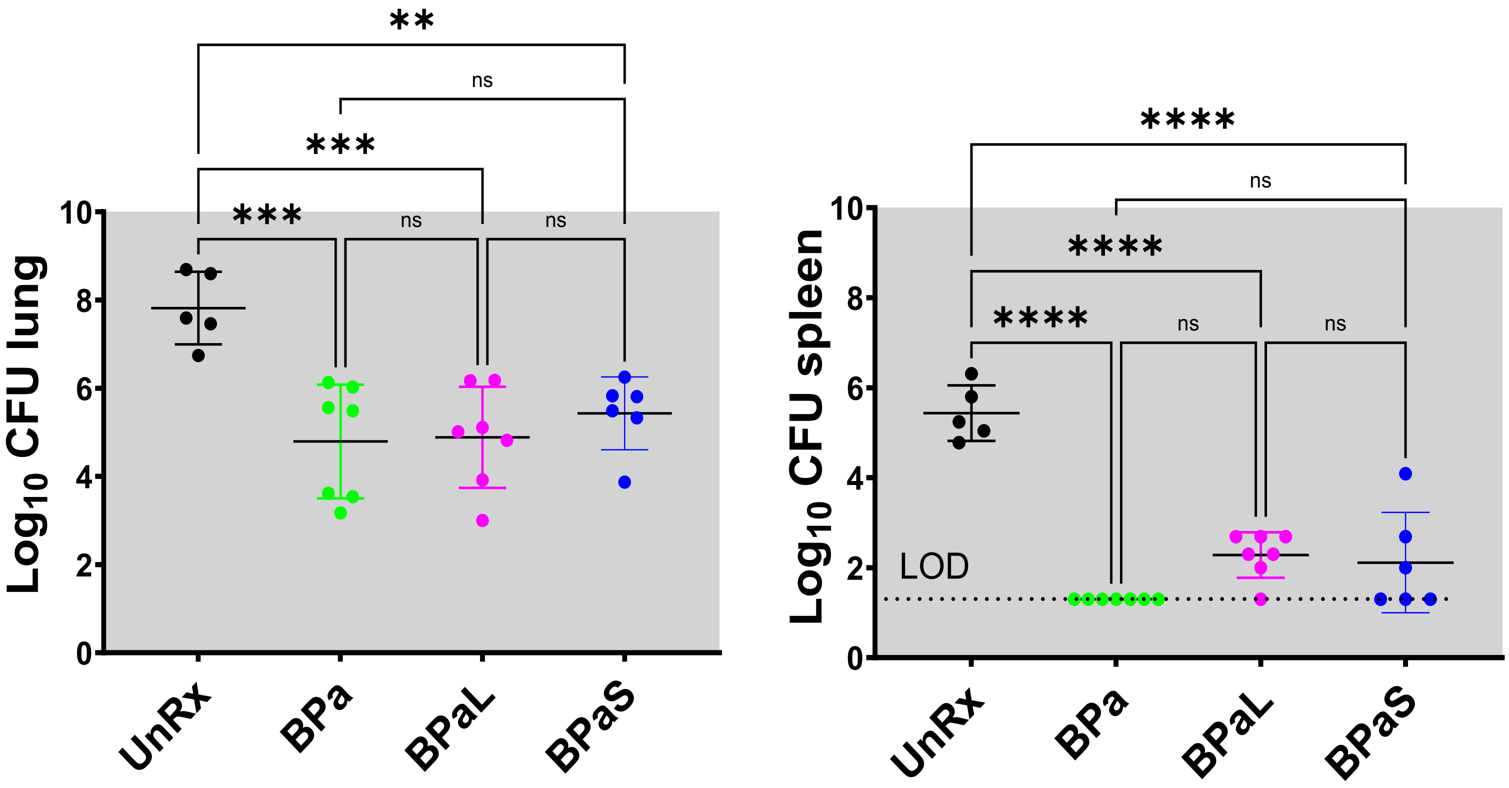


**Figure S3 (C3HeB/FeJ Study 3).** Bacterial burden (CFU) in Mtb infected C3HeB/FeJ mice treated with BPa, BPaL and BPaS regimen for 4 weeks. n = 7, LOD: limit of detection, $p < 0.05$

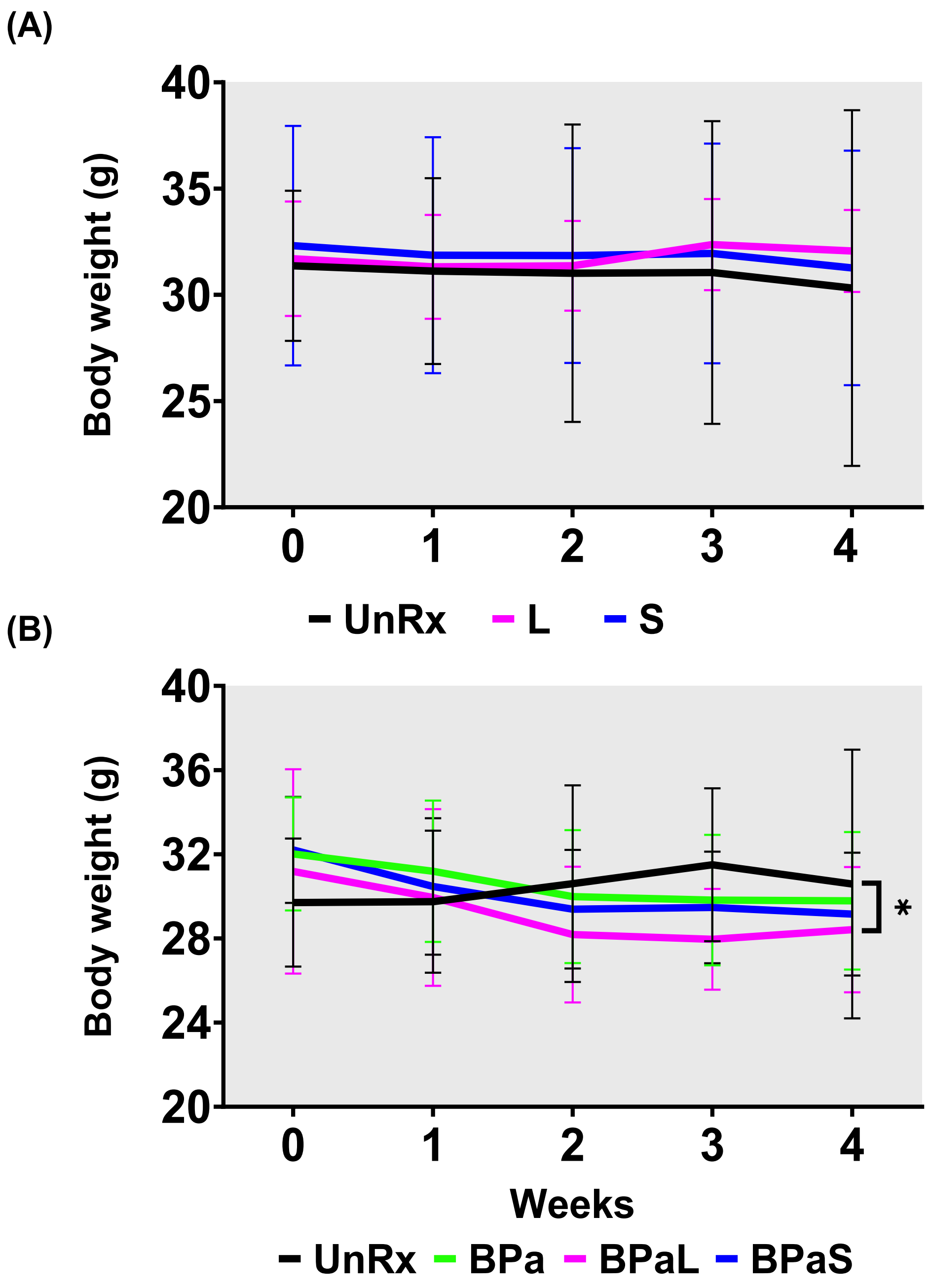


**Figure S4.** Change in the average body weight of Mtb infected C3HeB/FeJ mice during drug treatment. (A) represents the change during monotherapy of linezolid (L) and spectinamide 1599 (S) compared to untreated (UnRx) control while (B) represents the combined data from three independent studies during combination treatment with BPa, BPaL and BPaS. $p < 0.05$

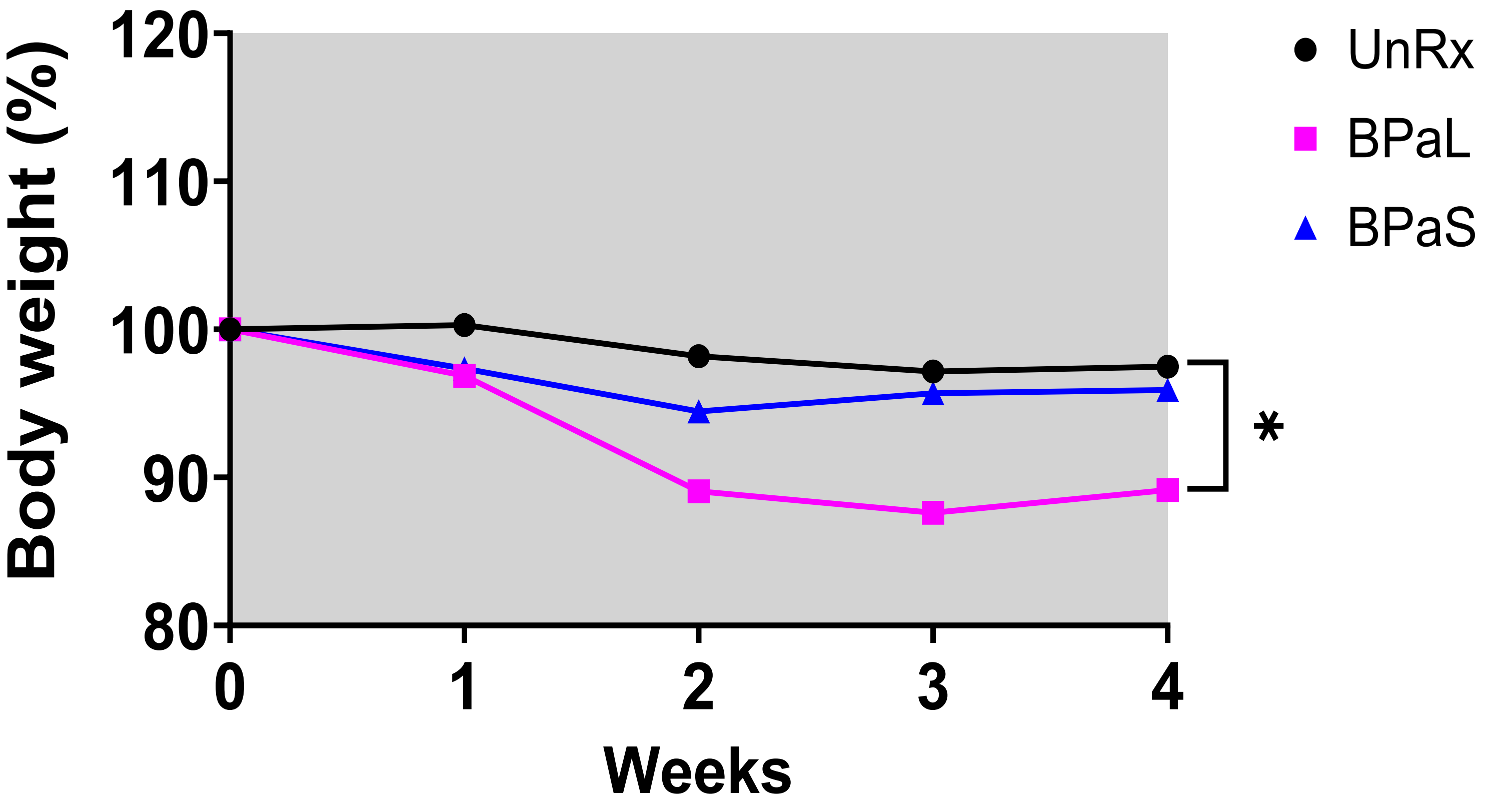


**Figure S5 (C3HeB/FeJ Study 1).** Change in the average body weight of Mtb infected C3HeB/FeJ mice during drug treatment. n = 7, $p < 0.05$

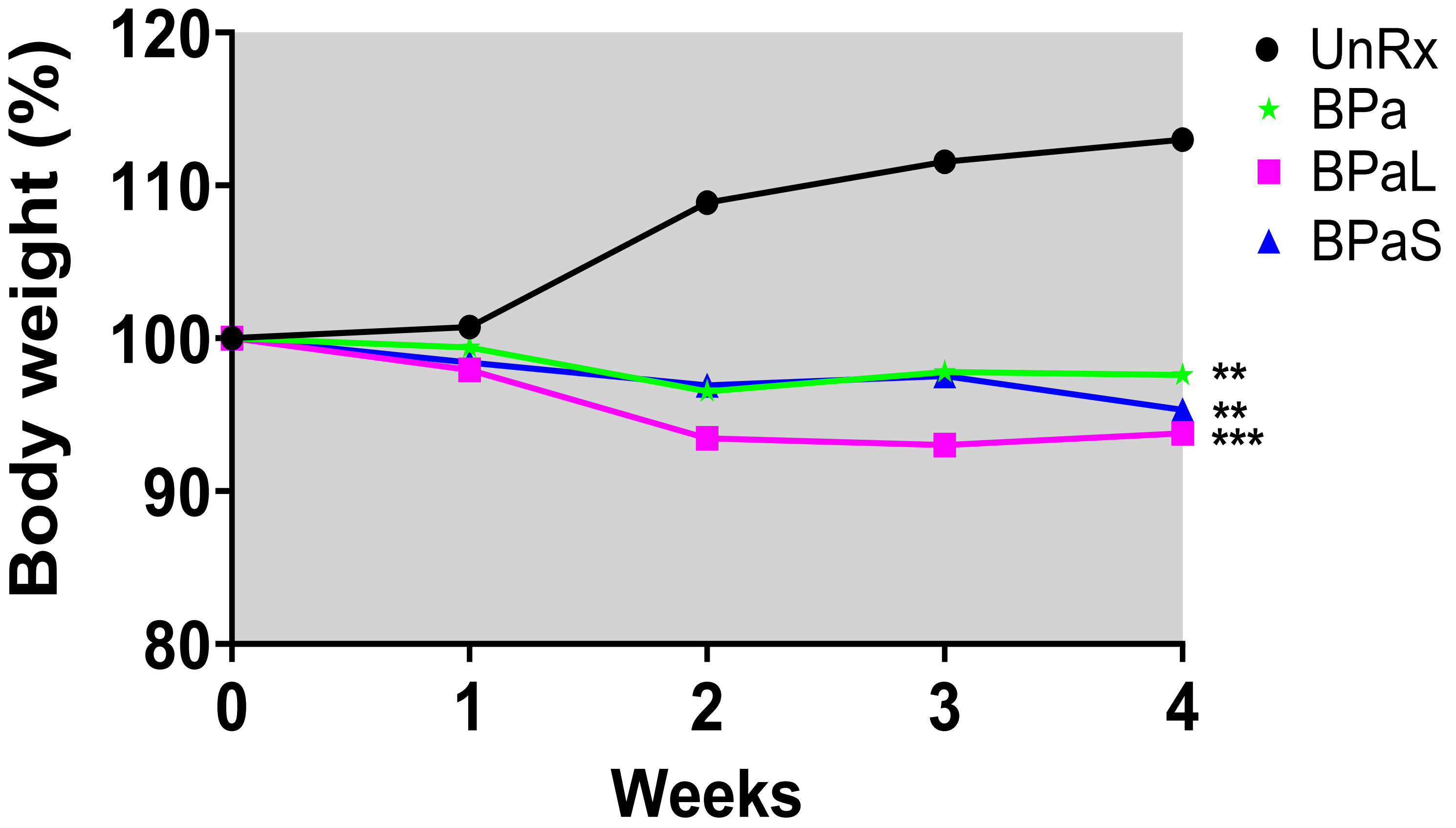


**Figure S6 (C3HeB/FeJ Study 2).** Change in the average body weight of Mtb infected C3HeB/FeJ mice during drug treatment. n = 3-9, $p < 0.05$

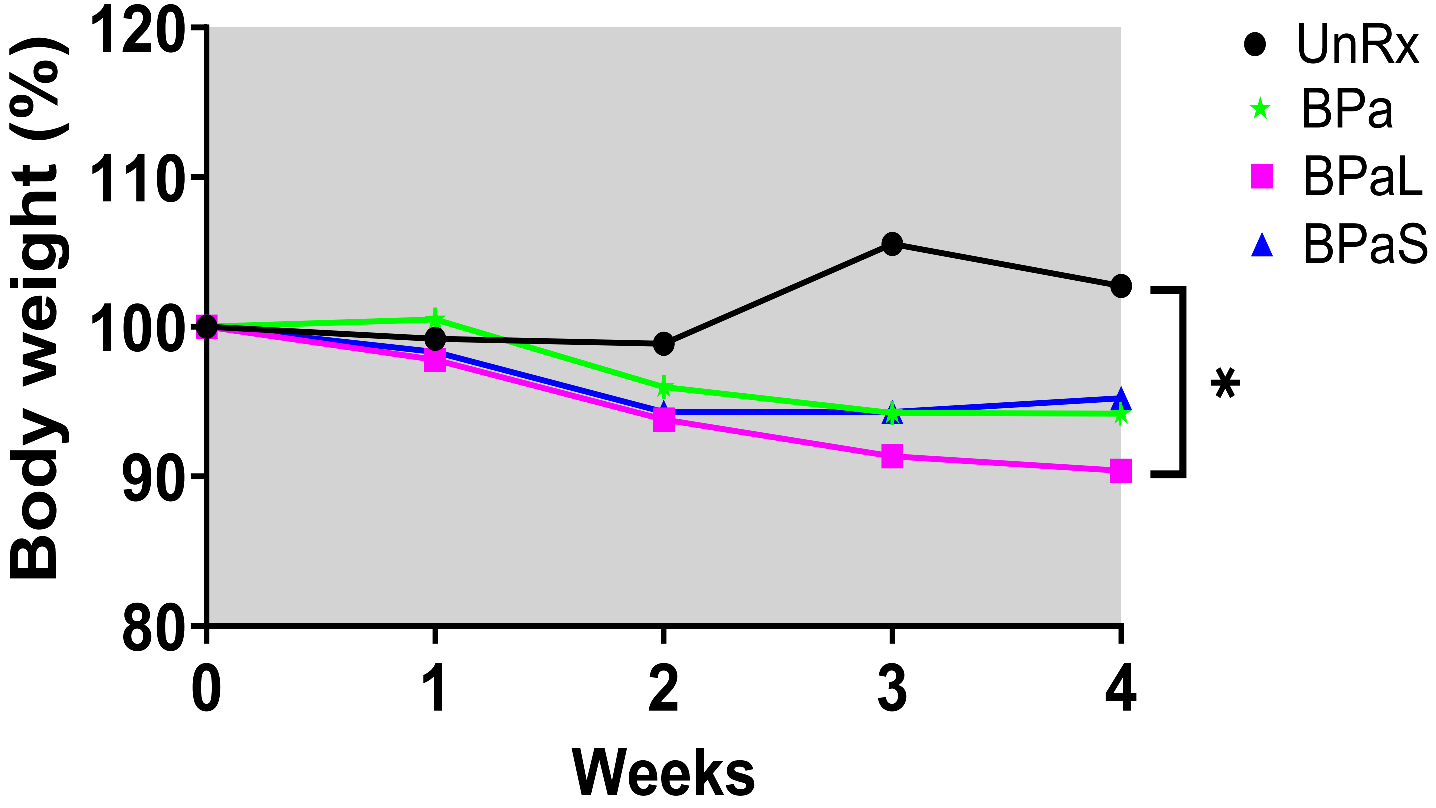


**Figure S7 (C3HeB/FeJ Study 3).** Change in the average body weight of Mtb infected C3HeB/FeJ mice during drug treatment. n = 7, $p < 0.05$

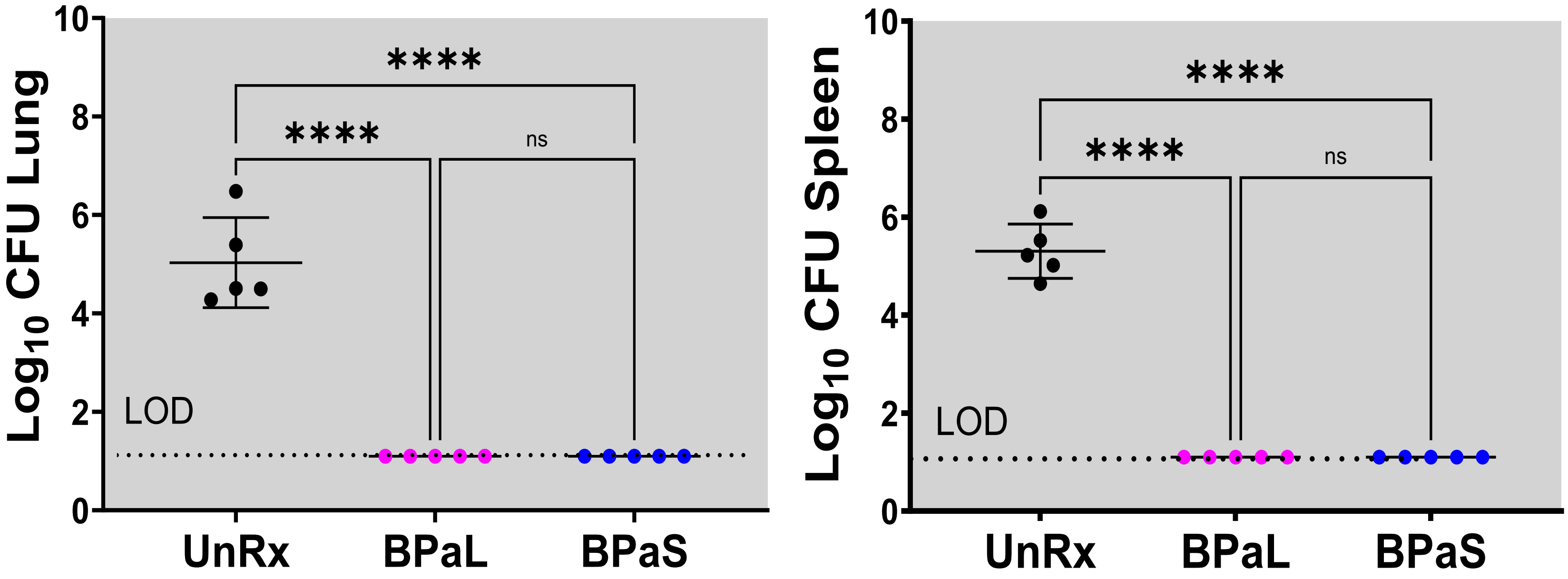


**Figure S8 (BALB/c Study 1).** Bacterial burden (CFU) in Mtb infected BALB/c mice treated with BPaL and BPaS regimen for 4 weeks. n = 5, LOD: limit of detection, $p < 0.05$

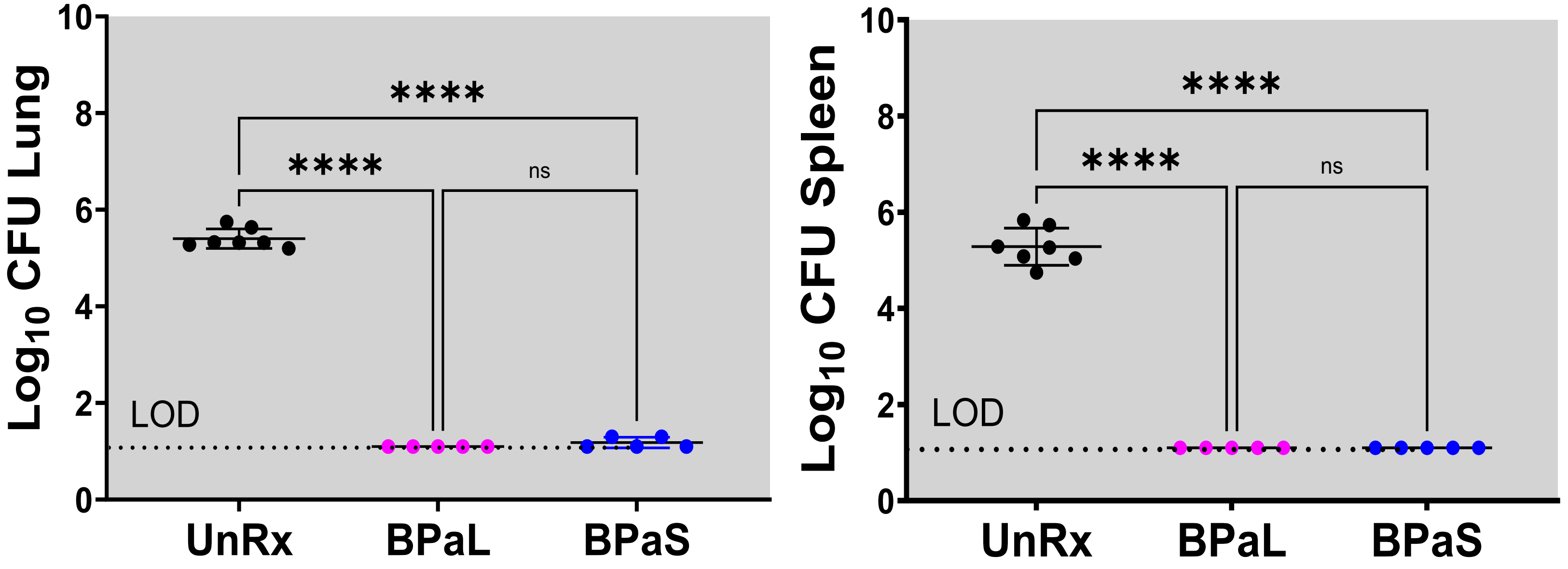


**Figure S9 (BALB/c Study 2).** Bacterial burden (CFU) in Mtb infected BALB/c mice treated with BPaL and BPaS regimen for 4 weeks. n = 7, LOD: limit of detection, $p < 0.05$

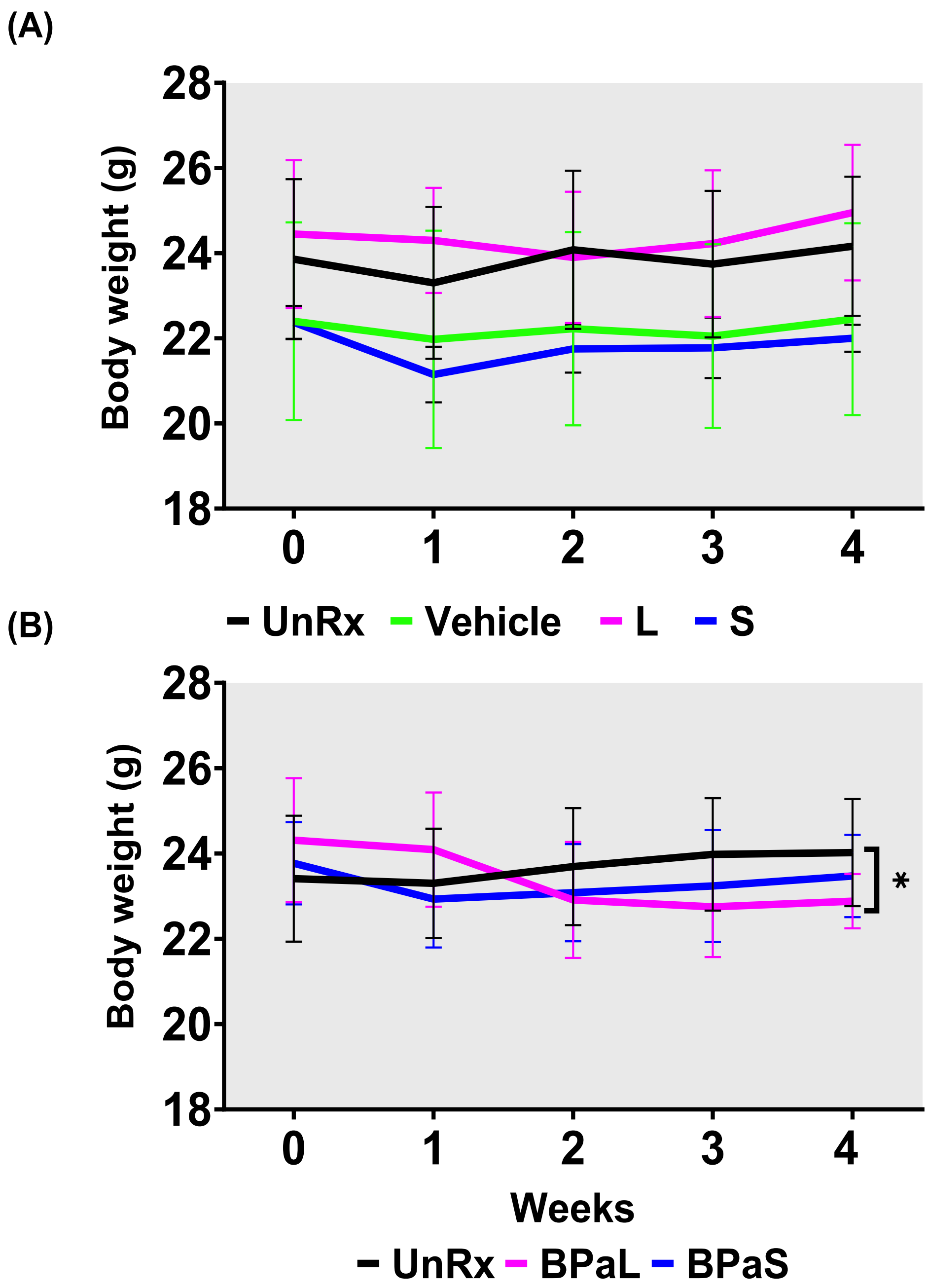


**Figure S10.** Change in the average body weight of Mtb infected BALB/c mice during drug treatment. (A) represents the change during monotherapy of linezolid (L) and spectinamide 1599 (S) compared to untreated (UnRx) and vehicle control while (B) represents the combined data from two independent studies during combination treatment with BPaL and BPaS. $p < 0.05$

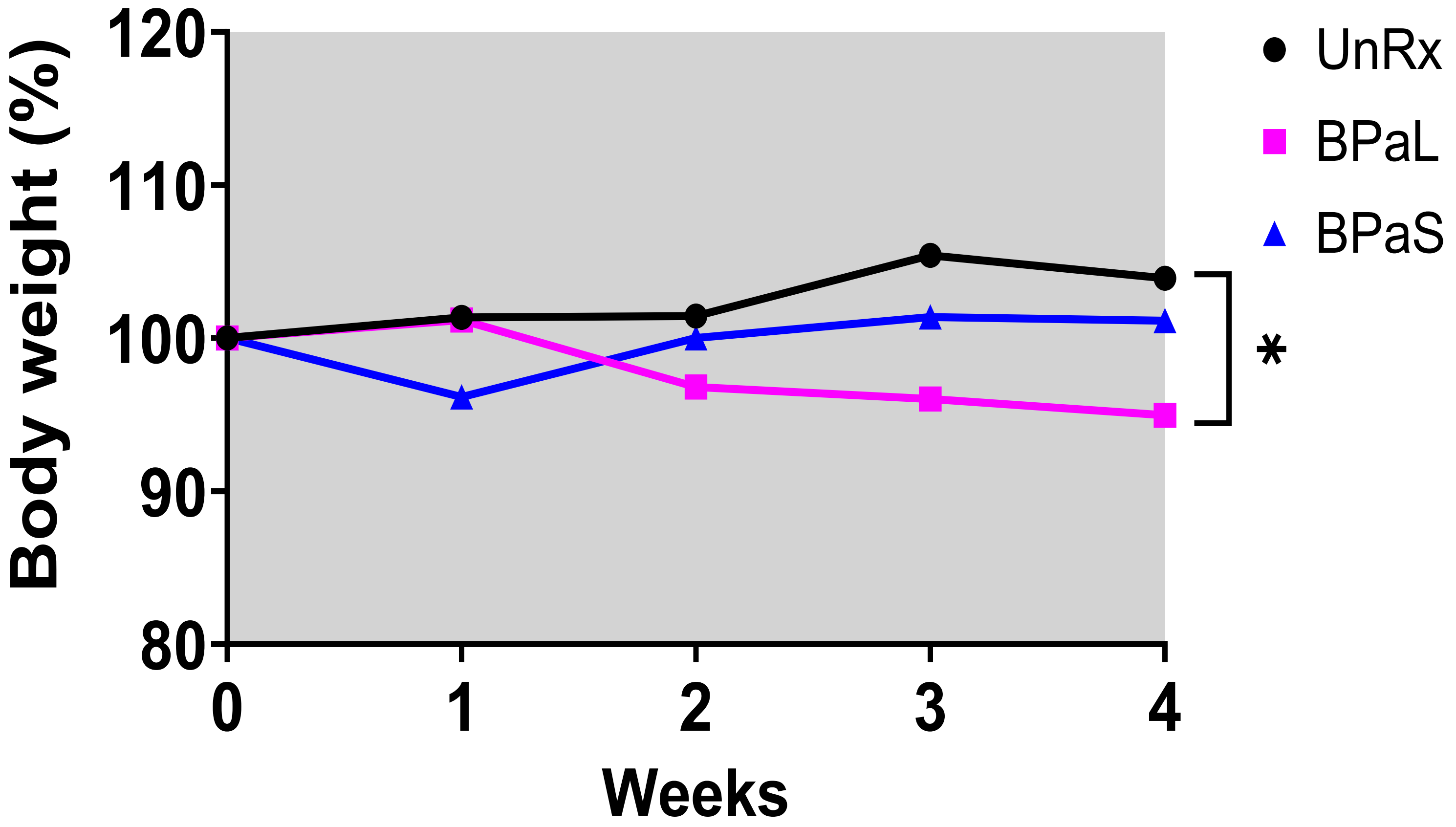


**Figure S11 (BALB/c Study 1).** Change in the average body weight of Mtb infected BALB/c mice during drug treatment. n = 5, $p < 0.05$

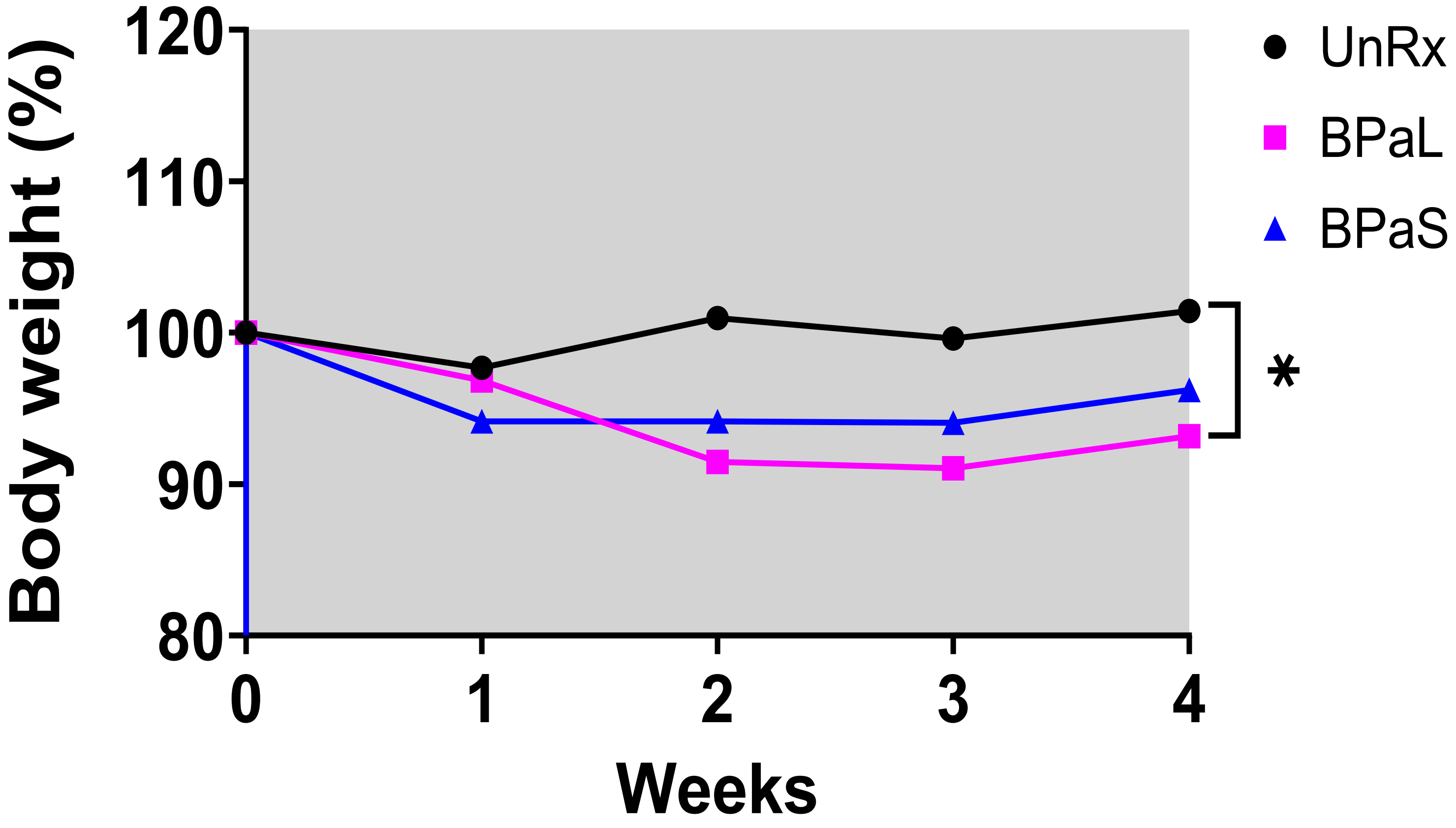


**Figure S12 (BALB/c Study 2).** Change in the average body weight of Mtb infected BALB/c mice during drug treatment. n = 7, $p < 0.05$

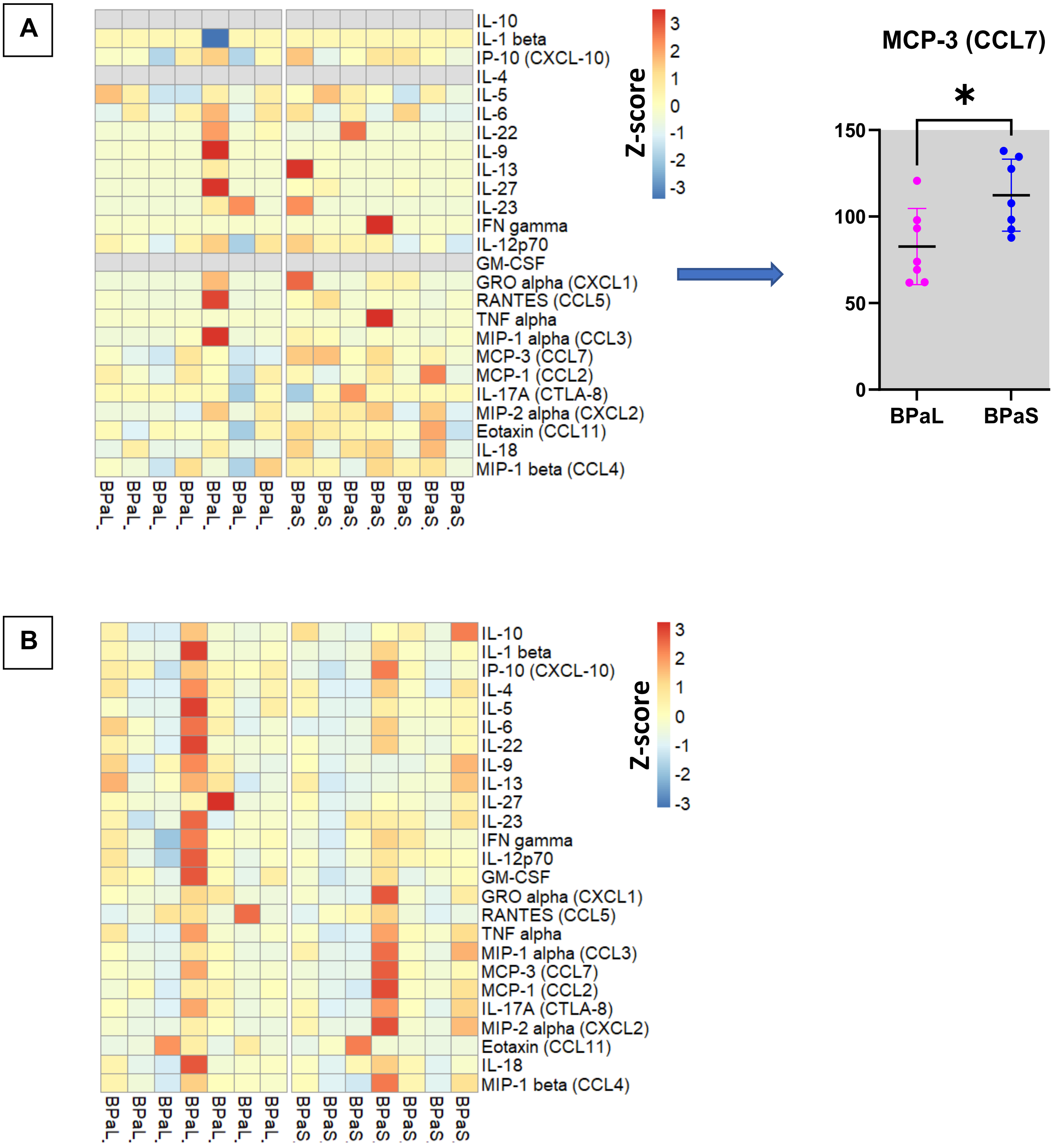


**Figure S13**. Change in the cytokines and chemokines profile in Mtb infected C3HeB/FeJ mice during drug treatment. (A) Plasma and (B) lung cytokine and chemokine contents in mice treated with BPaL or BPaS.

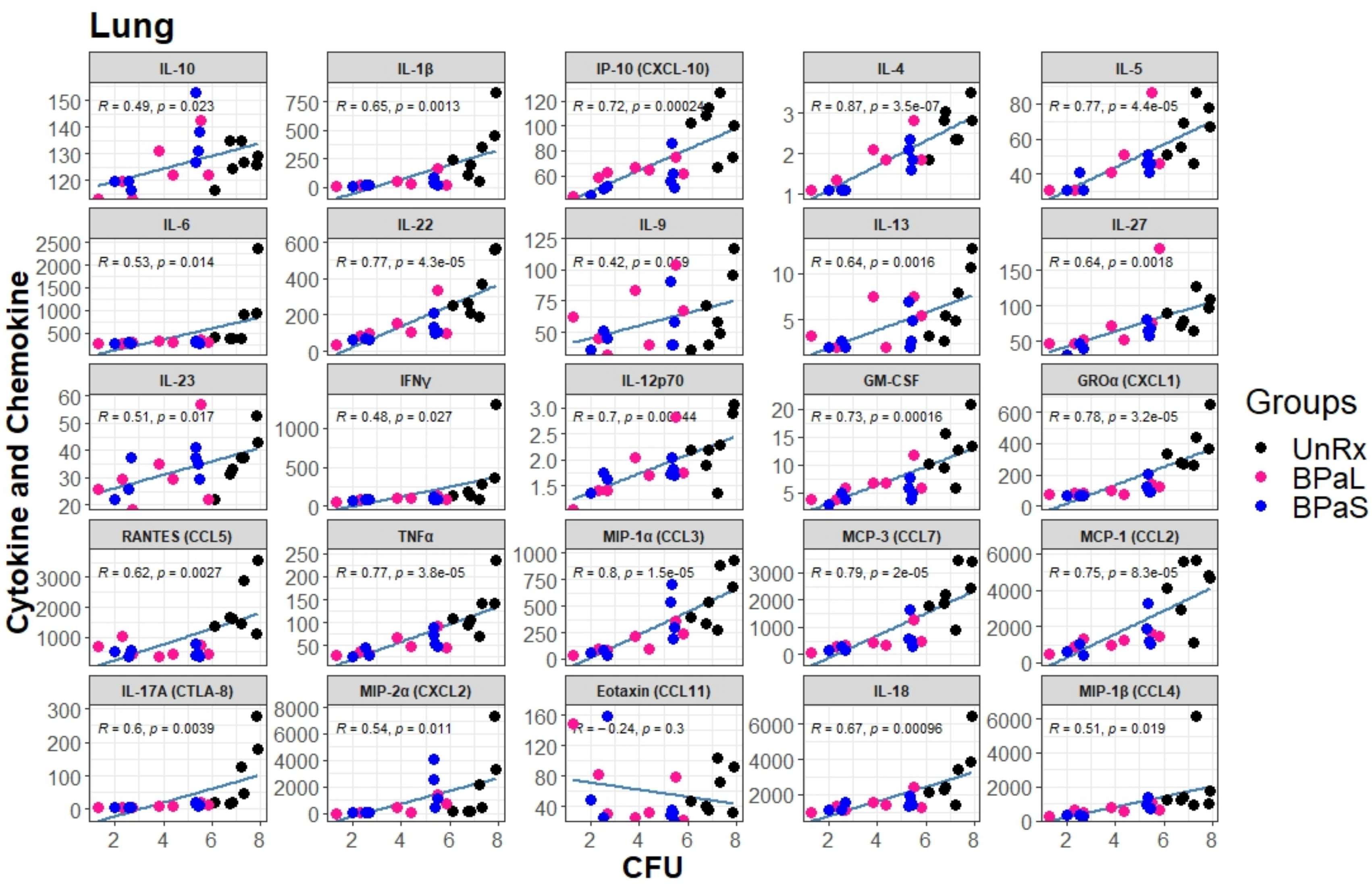


**Figure S14**. Spearman's correlation analysis between bacterial burden (CFU) and cytokine and chemokine profile in the lungs of Mtb infected C3HeB/FeJ mice treated with BPaL and BPaS regimen.

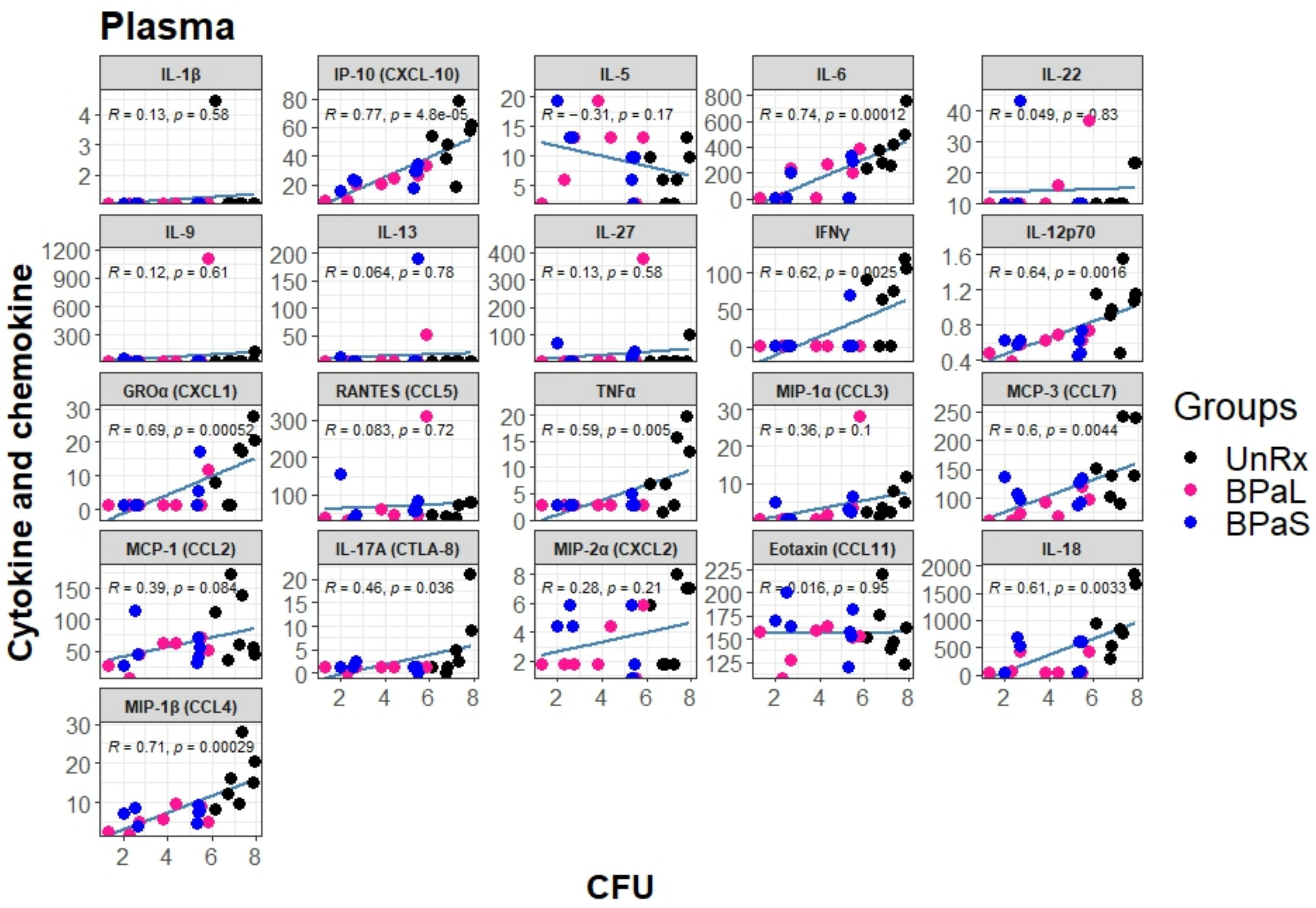


**Figure S15**. Spearman's correlation analysis between lung bacterial burden (CFU) and plasma cytokine and chemokine profile in Mtb infected C3HeB/FeJ mice treated with BPaL and BPaS regimen.

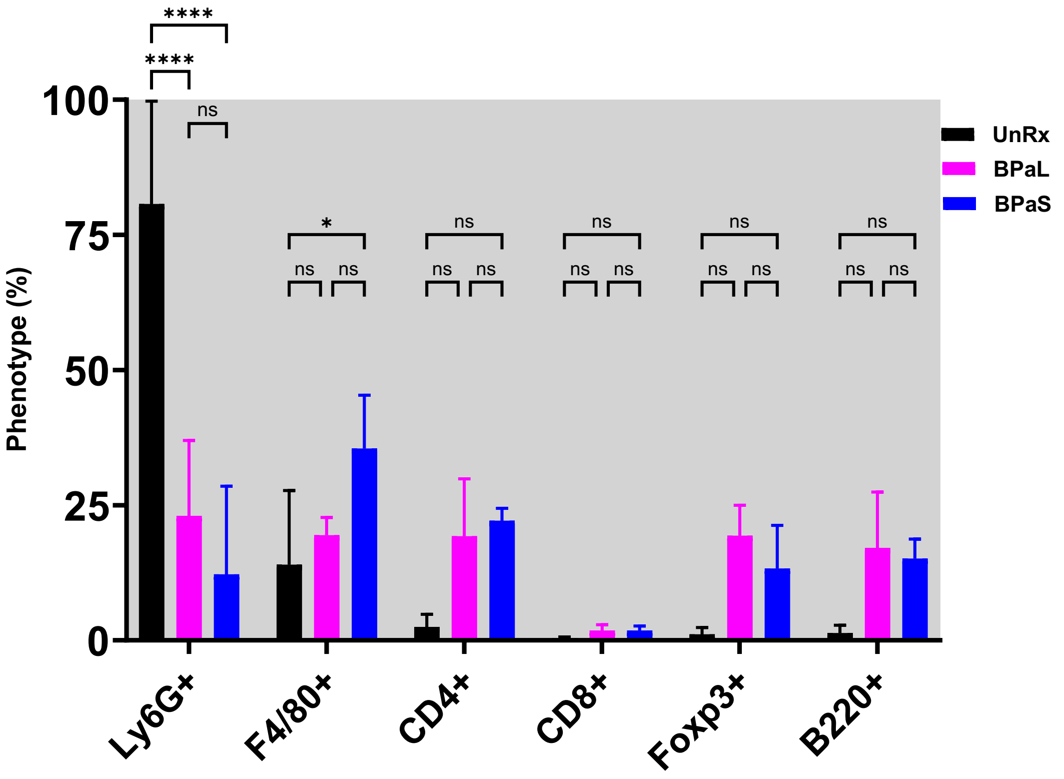


**Figure S16.** Immune cell populations in the lungs of Mtb infected C3HeB/FeJ TB model after 4 weeks therapy. Statistical significance was calculated using two-way ANOVA with Tukey's test for multiple comparisons and $p < 0.05$ was considered significant.

**Table S1**

| REAGENT or RESOURCE | SOURCE | IDENTIFIER |
|---|---|---|
| **Antibodies** | | |
| Anti-mouse LY6G PerCP | BioLegend | Cat# 127654; RRID: AB_11218876 |
| Anti-mouse CD14 PerCP Cy5.5 | Invitrogen | Cat# 120606; RRID: AB_493267 |
| Anti-mouse NKp46/CD335 PE | BioLegend | Cat# 137604; RRID: AB_2566163 |
| Anti-mouse B220/ CD45R PE-Cy7 | BioLegend | Cat# 103222; RRID: AB_2573837 |
| Anti-mouse CD8 FITC | BioLegend | Cat# 100706; RRID: AB_394458 |
| Anti-mouse CD34 PE-Dazzle 594 | BioLegend | Cat# 128616; RRID: AB_11219403 |
| Anti-mouse TER119 APC | BD Pharmingen | Cat# 561033; RRID: AB_10900980 |
| Anti-mouse γδ-TCR APC Fire 750 | BioLegend | Cat# 118129; RRID: AB_755986 |
| Anti-mouse LY6C Alexa Fluor 700 | BioLegend | Cat# 128024; RRID: AB_2869739 |
| Anti-mouse CD4 BV421 | BioLegend | Cat# 100544; RRID: AB_2562555 |
| Anti-mouse MHC-II BV480 | BD Biosciences | Cat# 566088; RRID: AB_2562612 |
| Anti-mouse CD11b Pacific Blue | BioLegend | Cat# 101224; RRID: AB_2565937 |
| Anti-mouse CD3e BV510 | BioLegend | Cat# 100353; RRID: AB_2563056 |
| Anti-mouse CD45 BV570 | BioLegend | Cat# 103136; RRID: AB_2814047 |
| Anti-mouse CD19 BV605 | BioLegend | Cat# 115540; RRID: AB_2563289 |
| Anti-mouse CCR2 BV711 | BD Biosciences | Cat# 747964; RRID: AB_2660295 |
| Anti-mouse CC11c BV785 | BioLegend | Cat# 117335; RRID: AB_2073247 |

**Table S2**

| Antibody | Specie | Type | Company | Catalogue # | Concentration | pH | Opal |
|---|---|---|---|---|---|---|---|
| CD8 | Rabbit | mAb | CST | D4W2Z | 1:400 | 6 | 480 |
| CD4 | Rat | mAb | Thermo Fisher | 4SM95 | 1:200 | 6 | 520 |
| B220 | Rat | mAb | BD Pharm | RA3-6B2 | 1:500 | 6 | 570 |
| FoxP3 | Rabbit | mAb | R&D | MAB8214 | 1:200 | 6 | 620 |
| Ly6G | Rabbit | mAb | CST | 87048 | 1:100 | 6 | 690 |
| F4/80 | Rabbit | mAb | CST | D4C8V | 1:100 | 6 | 780 |
